# Exogenous extracellular matrix proteins decrease cardiac fibroblast activation in stiffening microenvironment through CAPG

**DOI:** 10.1101/2021.06.05.447170

**Authors:** Xinming Wang, Valinteshley Pierre, Chao Liu, Subhadip Senapati, Paul S.-H. Park, Samuel E. Senyo

## Abstract

Controlling fibrosis is an essential part of regenerating the post-ischemic heart. In the post-ischemic heart, fibroblasts differentiate to myofibroblasts that produce collagen-rich matrix to physically stabilize the infarct area. Infarct models in adult mice result in permanent scarring unlike newborn animals which fully regenerate. Decellularized extracellular matrix (dECM) hydrogels derived from early-aged hearts have been shown to be a transplantable therapy that preserves heart function and stimulates cardiomyocyte proliferation and vascularization. In this study, we investigate the anti-fibrotic effects of injectable dECM hydrogels in a cardiac explant model in the context of age-associated tissue compliance. Treatments with adult and fetal dECM hydrogels were tested for molecular effects on cardiac fibroblast activation and fibrosis. Altered sensitivity of fibroblasts to the mechanosignaling of the remodeling microenvironment was evaluated by manipulating the native extracellular matrix in explants and also with elastomeric substrates in the presence of dECM hydrogels. The injectable fetal dECM hydrogel treatment decreases fibroblast activation and contractility and lowers the stiffness-mediated increases in fibroblast activation observed in stiffened explants. The anti-fibrotic effect of dECM hydrogel is most observable at highest stiffness. Experiments with primary cells on elastomeric substrates with dECM treatment support this phenomenon. Transcriptome analysis indicated that dECM hydrogels affect cytoskeleton related signaling including Macrophage capping protein (CAPG) and Leupaxin (LPXN). CAPG was down-regulated by the fetal dECM hydrogel. LPXN expression was decreased by stiffening the explants; however, this effect was reversed by dECM hydrogel treatment. Pharmacological disruption of cytoskeleton polymerization lowered fibroblast activation and CAPG levels. Knocking down CAPG expression with siRNA inhibited fibroblast activation and collagen deposition. Collectively, fibroblast activation is dependent on cooperative action of extracellular molecular signals and mechanosignaling by cytoskeletal integrity.

## INTRODUCTION

Cardiovascular disease is the leading cause of death worldwide. Up to 1 billion cardiomyocytes die after myocardial infarction (MI), or heart attack [1]. Neonatal mice and pigs can fully recover from a surgical MI model within a month [2]–[4]. The high cardiac regenerative capacity is lost with maturity towards adolescence and adulthood [5]. After an MI, necrotic tissue is permanently replaced with fibrotic extracellular matrix (ECM) which protects the heart from rupture by maintaining structural integrity. Nevertheless, the changing mechanical stress post-MI stimulates fibrotic deposition that extends beyond the acute injury zone. Excessive fibrosis results in reduced chamber compliance and increased ventricle wall stiffness which compromises cardiac function [6].

Fibroblasts regulate the extracellular matrix dynamics. After an MI, fibroblasts in the infarct area proliferate and differentiate to the transient myofibroblast state [7]. Myofibroblasts shift the molecular profile of extracellular matrix and increase matrix protein deposition to stabilize the infarct area. Collagen I and fibronectin are the major extracellular fiber proteins produced by myofibroblast. Collagen is crosslinked by lysyl oxidase (LOX) enzyme which effectively maintains the structural stability of extracellular matrix. Myofibroblasts are further characterized as expressing α-smooth muscle actin (α-SMA) and elevated traction force to then increase the tensile stress of the newly synthesized matrix.

Fibroblasts are sensitive to the mechanical properties of the microenvironment. For proof-of-concept, cardiac fibroblasts cultured on stiff elastomeric substrates (elastic modulus of 8kPa) were shown to express α-SMA protein and synthesize more collagen than on soft substrates (3kPa) [8], [9]. Transforming growth factor-β1 (TGF-β) induced differentiation of lung fibroblasts to myofibroblasts was enhanced by increasing the substrate stiffness [10]. Yes-associated protein 1 (YAP) and Rho-associated protein kinase (ROCK) have been reported to mediate the mechanosensitivity in many cells including fibroblasts [9], [11], [12]. We recently demonstrated that transplanting decellularized ECM (dECM) induced a pro-proliferative phenotype in cardiomyocytes in mice and explants by mechanosensitive signaling pathways including YAP. Here we extend that work to investigate the phenomenon of decreased fibrosis with softened microenvironment stiffness and dECM treatments [13].

Because Rho kinase is an important node in fibroblast activation and fibrosis, proteins that regulate Rho signaling pathway and cytoskeleton organization may be relevant to fibroblast activation attenuation by dECM [14]–[16]. Macrophage capping protein (CAPG) is an actin regulatory protein that has not been evaluated in cardiac fibroblasts. Inhibiting CAPG expression lowers macrophage ruffling and cancer metastasis[17], [18]. Increasing CAPG expression promotes smooth muscle cell migration under hypoxia and 3T3 cell ruffling induced by platelet-derived growth factor [19], [20]. Leupaxin (LPXN) is a member of paxillin family of focal adhesion proteins. LPXN has been reported to regulate cell adhesion and migration [21]–[23]. LPXN suppresses the tyrosine phosphorylation of paxillin which influences various pathways [24], [25]. CAPG and LPXN were investigated for a role in dECM regenerative signaling.

Decellularized extracellular matrix (dECM) hydrogel is a natural biomaterial initially employed for tissue engineering. The decellularized organ is liquefied by enzymatic digestion before injecting into tissue. Porcine heart derived dECM hydrogel has been clinically tested in heart failure patients for safety [26]. In preclinical experiments, delivering dECM hydrogel derived from regenerative hearts into the injured adult rodent hearts prevents the spreading of infarct area, protects heart function, and stimulates vascularization [27], [28]. We have shown that the dECM hydrogel induced pro-proliferative phenotype in cardiomyocytes can be increased by softening the microenvironment stiffness in heart explants [13]. The mechanism of dECM hydrogel induced heart regeneration has not been fully understood. The presence of extracellular proteins, like TGF-β1 and agrin, in the dECM hydrogel may modulate the cardiac post-injury response.

BAPN has been used *in vivo* to decrease the stiffness of hearts [13], [29]. BAPN modulates extracellular matrix stiffness by inhibiting LOX activity and thus reduces collagen crosslinking. Ribose increases extracellular matrix stiffness by crosslinking proteins through glycation. Ribose has been shown to increase the stiffness of collagen gels and cardiac explants [13], [30]. There is no evidence to suggest that BAPN and ribose directly affect fibroblast activation. We previously demonstrated that BAPN and ribose treatment do not alter cardiomyocyte cell cycle activity and morphology in heart explants [13].

We previously demonstrated that cardiomyocyte cell cycle activity induced by fetal dECM is mechanosensitive [13]. Here, we hypothesize that intersecting signaling from decellularized matrix transplantation and mechanical stimuli lowers fibroblast activation resulting in reduced collagen deposition. To test the hypothesis, the stiffness of extracellular matrix in explants was modulated by BAPN and ribose. Exogenous dECM hydrogel was investigated for a role in fibroblast activation and fibrosis in an explant model of the left ventricle with tuned microenvironment stiffness. We examined the cooperative-effect of exogenous dECM hydrogel and the microenvironment stiffness on fibroblast activation in 3D-explants and on 2D-polymer substrates. Explant transcriptomes were analyzed to determine how dECM hydrogel influenced fibroblast activation which indicated cytoskeletal regulation. Cytoskeleton polymerization and CAPG expression were modulated and establish a role for CAPG in dECM hydrogel regulated fibroblast activation.

## METHODS

### P1 mice myocardial infarction model

Animal procedures were reviewed and approved by the Institutional Animal Care and Use Committee at the Case Western Reserve University according to the guidelines and regulations described in the Guide for the Care and Use of Laboratory Animals (National Academies Press, 2011). All animals were maintained and housed under specific pathogen-free conditions at our animal facility accredited by the Association for Assessment and Accreditation of Laboratory Animal Care, International (AAALAC) at Case Western Reserve University.

Surgical MI protocol performed on neonatal mice was adopted from published work[31]. Briefly, anesthesia was induced by hypothermia on day 1 mice. A cut was made on the fourth intercostal space to visualize the heart. Permanent MI was induced by ligating coronary artery with a 10-0 nylon suture. 20μg adult or fetal dECM in 2μl PBS or 2μl PBS (negative control) were injected into myocardium using 10μl Hamilton syringes, 1 injection below and 1 above the ligation site. Chest cavity was closed by suturing ribs and muscles, and skin was closed using skin glue. Mice were euthanized 1 week after surgery for histology.

### dECM preparation

Gelable dECM was prepared following published protocols [13]. Briefly, diced porcine adult or fetal hearts were decellularized by sodium dodecyl sulfate and Triton X-100 washing. After extensively washing in water, decellularized extracellular matrix (dECM) was lyophilized, cryo-pulverized, and enzymatically digested. The digested dECM was then adjusted to a neutral pH and stored at −20°C.

### Ventricle explant culture

Hearts harvested from day 1 mice were washed in cold 1 x PBS. Atrial was removed and ventricle was cut to approximately 1mm^3^ pieces using surgical scissors. Two microliters of adult or fetal dECM were injected into explants using 10μl Hamilton syringes with 23G needles before plating on 48-well plates coated with 1.5mg/ml collagen I (Corning) gel. Explants were incubated at 37°C for 2h to induce dECM polymerization and explant-collagen gel attachment. Explants were cultured in explant culture media [1% fetal bovine serum, 2mM L-glutamine, 1 x insulin-transferrin-selenium (Gibco, ITS-G 100X) in M199] for 6 days with 10μM BrdU labeling for 3 days before fixation. Paraformaldehyde (4%) was used for fixation. In separate experiments, explants were cultured in explant culture media supplemented with 5mM ribose or 0.2mM BAPN for 6 days from plating in combination with dECM treatment conditions. Atomic force microscopy was used to measure the elastic modulus of decellularized explants [13], [32].

In order to regulate cytoskeleton polymerization, explants were cultured in explant culture media supplemented with Jasplakinolide (Cayman), Latrunculin A (Cayman), Y27632 (Chemdea), PF573228 (Selleck) for 3 days; no BrdU labeling applied.

### Primary fibroblast isolation and culture

Primary rat ventricular fibroblasts were isolated from day 1 Sprague Dawley rats purchased from Charles River Laboratories US. Cardiac cells were dissociated from diced ventricular tissues using the Neonatal Heart Dissociation Kit, mouse and rat (Miltenyi Biotec). Cardiomyocytes and non-cardiomyocytes were separated by Percoll gradient centrifugation. Non-cardiomyocytes were plated on 75cm^2^ tissue culture flasks in 10% FBS DMEM and cultured in 5% FBS DMEM until reaching 80% confluent to then passage. After 2 passages, fibroblasts were plated in cell culture at a density of 26,000 cells/cm^2^. Fibroblasts were maintained in plating media overnight before culturing in non-serum DMEM containing 100μg/ml dECM for 1 day. Fibroblasts were then fixed in 4% PFA.

### RNA interference

Primary fibroblasts were cultured in antibiotics supplemented DMEM without serum. CAPG siRNA (ThermoFisher, Silencer, assay ID 60569) and Lipofectamine RNAiMAX (ThermoFisher) were prepared following the manufacturer protocol and added to cell culture media. The final siRNA concentration was 200nM. Control cells were transfected with negative control siRNA (ThermoFisher). Cells were incubated with siRNA for 2 days and then treated with 100ug/ml solubilized fetal dECM or 10ng/ml transforming growth factor β (TGF-β) for 24h. Cells were then fixed.

### Histology and immunostaining

Explants and hearts were embedded in paraffin or optimal cutting temperature compound and sectioned to 4 or 5μm slides. Masson’s trichrome staining was used to show fibrotic tissue. Tissue sections or primary fibroblasts cultured on polyacrylamide substrates were immunostained for α-SMA (Cell signaling technology), platelet-derived growth factor receptor alpha (pdgfr-α) (Abclonal), CAPG (ProteinTech), LPXN (ProteinTech), and vimentin (Abcam) antibodies, and counterstained for DAPI. Alexa Fluor conjugated secondary antibodies (ThermoFisher Scientific) were used for detection.

Samples were imaged within 48 hours after staining. Four pictures were randomly taken for the fibroblasts cultured on polyacrylamide, and within the area 200μm from the edge of explant sections. Explant exposed surfaces exhibit excess collagen deposition mimicking the infarct border region in hearts. For in vivo experiments, six pictures were randomly taken in the infarct and border zone from heart sections.

### mRNA sequencing and analysis

Total RNA was extracted from explants using RNA extraction kit (QIAGEN). RNA degradation was examined by agarose gel electrophoresis, purity examined by nanodrop (Thermo Fisher), quantify examined by Qubit (Thermo Fisher), and integrity examined by Agilent 2100 (Agilent). The mRNA was enriched by oligo beads, fragmented randomly, and used for cDNA synthesis. After purification and enrichment, cDNA was sequenced using HiSeq systems (Illumina). Data mapping performed by TopHat2 with mismatch parameter set to 2 and other parameters set to default. The gene counting data was transferred to reads per kilobase per million mapped reads for differential expression analysis. Gene ontology (GO) and protein-protein interaction were analyzed by STRING [33].

### Western blot

Proteins were extracted from explants using cell extraction buffer (Thermo Fisher). Equivalent protein lysate (20μg) was loaded in each lane of 4-20% polyacrylamide gel (Bio-Rad) for electrophoresis. Proteins were transferred to 0.45μm nitrocellulose membrane by semi-dry transferring. Membranes were blocked in 4% non-fat milk, followed by 2-hour incubation with 1:1000 diluted anti-LPXN, anti-CAPG, 1:60 diluted anti-collagen 1α1 (Developmental Studies Hybridoma Bank), and 1:20 000 diluted anti-GAPDH antibodies (Proteintech). Membranes were then incubated with 1:10,000 diluted HRP-conjugated secondary antibodies (Cell Signaling) for detection. Membranes were stripped in mild stripping buffer before re-probing.

### Statistical analysis

Data were shown as mean ± standard deviation. One-way ANOVA or Two-way ANOVA analyses was performed for comparing three or more groups. Tukey’s test was used for multiple comparisons. Confidential interval was 95%. T-test was used for comparing two groups. Confidence level was set to 95%. For each treatment in explant study, the experiment was repeated three times from different litters (biological replicates), with three to four explants per experiment (technical replicates). The total amount of cells analyzed in each treatment of the explant study is within 400 to 700. For isolated primary fibroblasts, three biological replicates with four technical replicates were used. The total amount of cells counted is within 400 to 1000.

## RESULTS

### dECM treatment reduces collagen deposition 7-days post MI in P1 mice

P1 neonatal mice hearts can fully regenerate in 3-weeks post-heart injury. However, at 7-days postsurgery, some scarring is observed in P1 mice. Investigating cardiac post-injury response in neonatal mice may reveal the mechanism of complete heart regeneration. Thus, a P1 neonatal mice MI model was used for determining the therapeutic efficacy of dECM treatments on heart injury.

*In vivo* experiments indicated dECM treatment lowered collagen deposition in P1 mice hearts a 1-week post-MI. Fibrosis and collagen deposition were examined by Masson’s trichrome staining on MI-control (Fig. 1A), fetal dECM (Fig. 1B), and adult dECM (Fig. 1C) treated hearts. Fetal dECM treated hearts showed significantly reduced collagen-rich area compared MI-control (Fig. 1D). Adult dECM treated hearts had a trend of lowering collagen deposition, but the change was not statistically significant.

**Figure 1.**
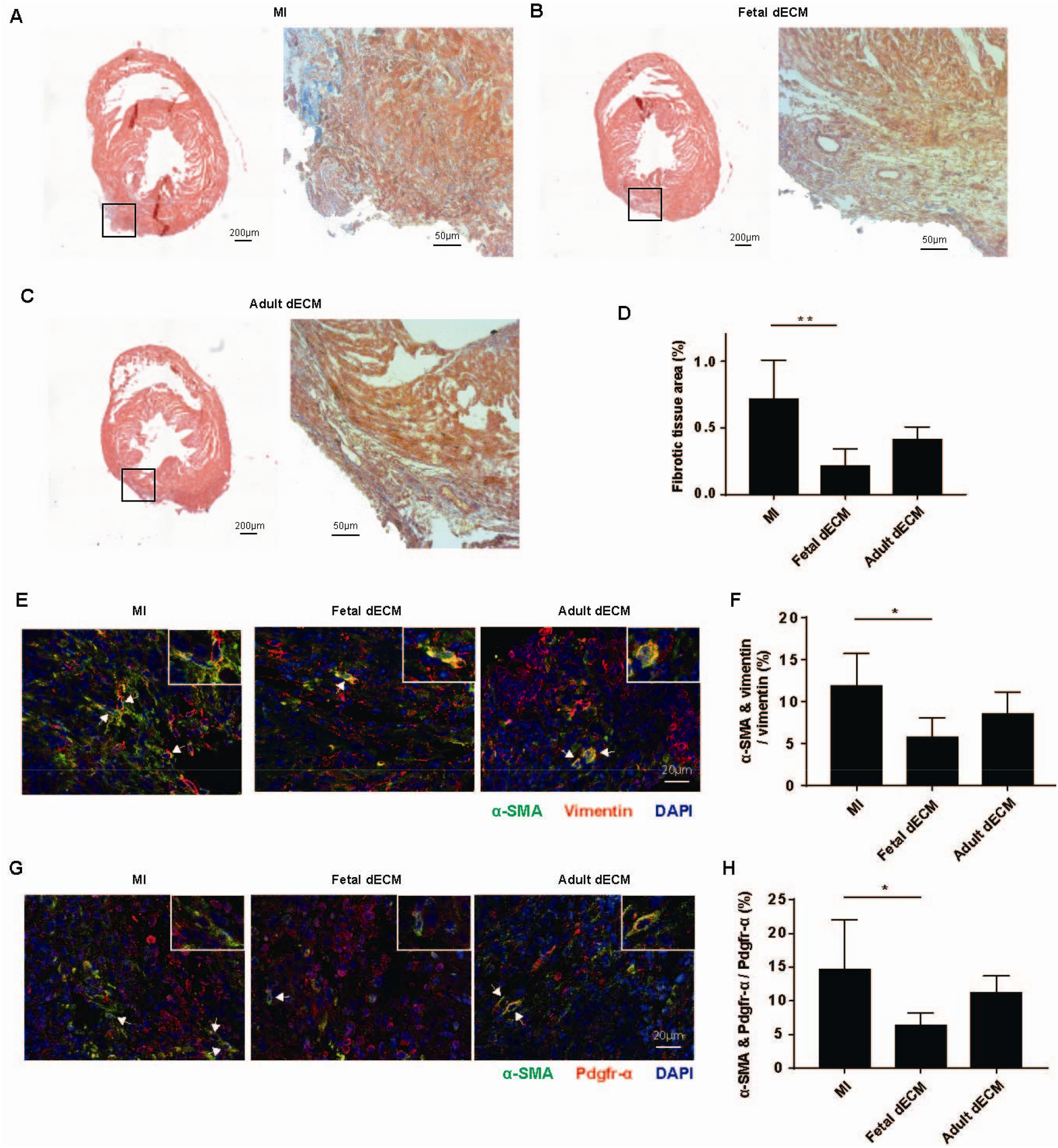
Fetal dECM reduces collagen deposition and fibroblast activation in P1 neonatal mice 7-days post-myocardial infarction. Collagen deposition was examined by Masson’s trichrome staining in (A) MI only, (B) fetal dECM injection, and (C) adult dECM injection hearts on day-7 post-MI. (D) The area of fibrotic tissue was significantly reduced by fetal dECM treatment but not by adult dECM. (E) Immunostaining of α-SMA and vimentin indicated (F) a lowered fibroblast activation in fetal dECM treated hearts. A similar result was observed in (G) α-SMA and pdgfr-α positive cells. (H) Fetal dECM reduced fibroblast activation. Adult dECM, however, did not significantly change fibroblast activation compared to MI hearts. (n=5 mice per group from 2 litters, one-way ANOVA and Tukey’s test, *p<0.05, **p<0.01, data presented as mean ± standard deviation. White arrow indicates α-SMA and vimentin / pdgfr-α doublepositive cell.)

Fibroblast activation was examined by immunostaining for α-SMA expression in vimentin (Fig. 1E) and pdgfr-α (Fig.1G) positive cells. Because of the heterogeneity of cardiac fibroblasts, there is no specific marker for cardiac fibroblasts [34]–[36]. We used two markers to identify the fibroblasts with the expectation that each of the markers labels a subtype of cardiac fibroblasts. In our experiment, fetal dECM treatment decreased the ratio of α-SMA positive cells in both vimentin (Fig. 1F) and pdgfr-α (Fig. 1H) positive cells compared to MI-control. Adult dECM, however, did not change α-SMA positive cells ratio relative to MI-control. The results indicate that fetal dECM, but not adult dECM, lowers fibrosis and fibroblast activation in P1 mice hearts on day-7 post-MI.

### Fetal dECM hydrogel reduces collagen deposition and fibroblast activation in P1 mice heart explants

We recently demonstrated that the neonatal regeneration model can be recapitulated with highly regenerative (early age) mice hearts explants in the dish to study post-injury response to transplanted dECM (porcine). Explants cut to 1mm cubes exhibit increased collagen deposition from the surface to a depth of approximately 200 microns that can be altered by age-sourced dECM [13]. Here, day 1 explants injected with fetal and adult dECM were stained for collagen deposition by Masson’s trichrome stain (Fig. 2A). In day 1 explants, fetal dECM hydrogel lowered the collagen border area compared to adult dECM hydrogel and no dECM treatment (Fig. 2B). In contrast, adult dECM hydrogel did not significantly change the fibrotic area. The results demonstrate that fetal dECM suppresses cardiac fibrosis in high regenerative heart explants.

**Figure 2.**
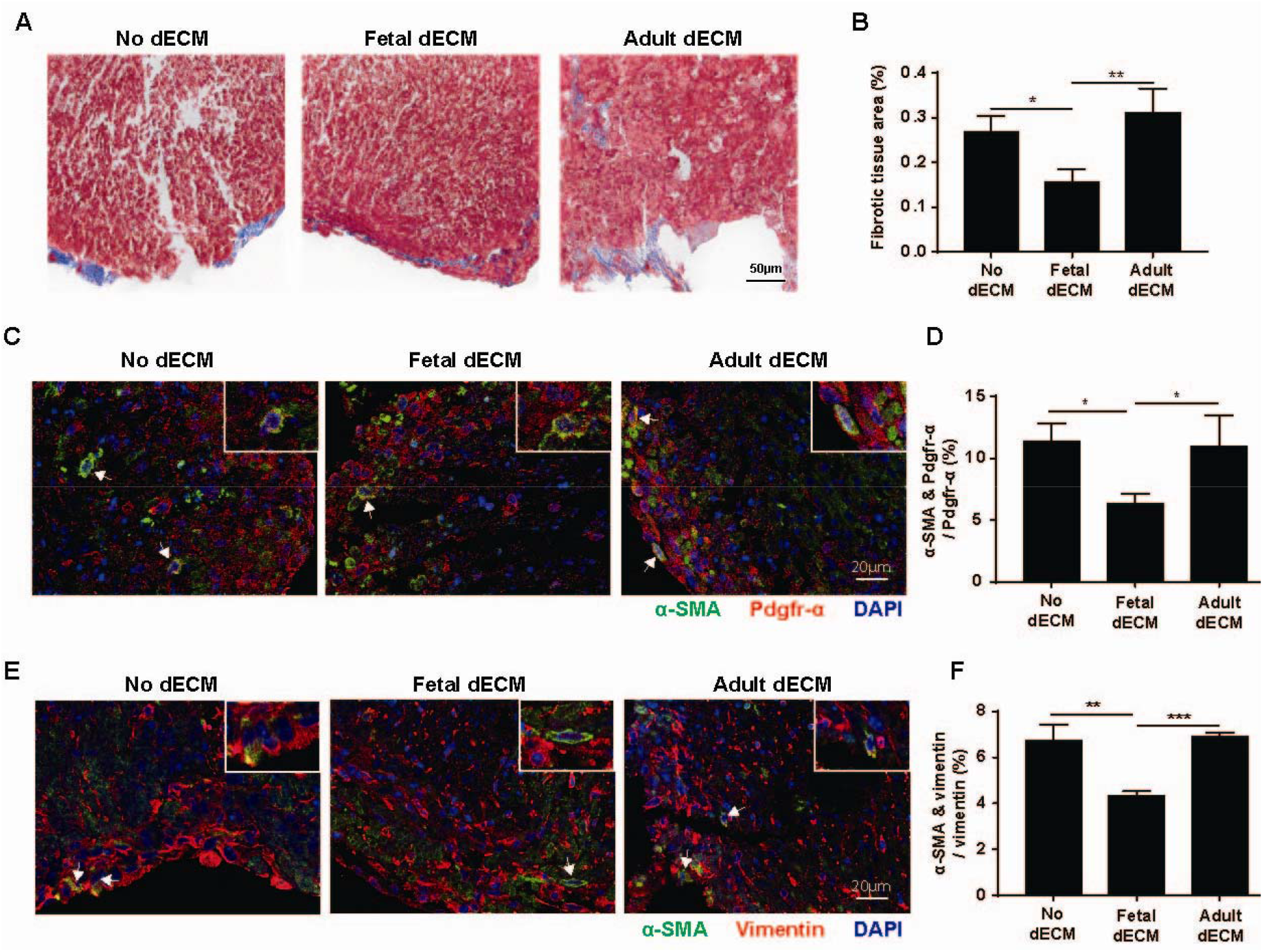
Fetal dECM hydrogel decreases collagen deposition and fibroblast activation in day 1 mice ventricle explants. (A) Collagen deposition in day 1 ventricle explants was stained by Masson’s trichrome staining. (B) Fetal dECM decreased collagen deposition in comparison to adult dECM and the control in day 1 explants. Adult dECM did not alter collagen deposition. (C) Fibroblasts activation was examined by immunostaining for α-SMA and pdgfr-α. (D) Fetal dECM lowered the ratio of α-SMA positive fibroblasts compared to the other treatments. (E) Day 1 explants were also stained for α-SMA and vimentin. (F) Fetal dECM reduced the number of α-SMA positive fibroblasts compared to control and adult dECM. (n=3 biological replicates, 3 explants in each biological replicate, one-way ANOVA and Tukey’s test, *p<0.05, **p<0.01, ***p<0.001, ****p<0.0001, data presented as mean ± standard deviation. White arrow indicates α-SMA and vimentin double-positive cell. Separate channels of a representative image in supplement figure 2A.)

The fibroblast activation was modulated by dECM treatments in explants. Fetal dECM treated day 1 explants had less activated fibroblasts compared to the other groups as indicated by immunostaining for α-SMA and pdgfr-α (Fig. 2C, D). A similar result was observed probing for α-SMA and vimentin positive cells (Fig. 2E, F). Adult dECM did not significantly change activated fibroblasts ratio compared to control. In a gel contraction assay, solubilized fetal dECM treatment also reduced the contractility of fibroblast (3T3) cell line embedded in a collagen gel (Supplement fig. 1A, B). Fibroblast activation was also examined by flow cytometry for α-SMA positive cell numbers (Supplement fig. 1C). Fetal dECM lowered α-SMA positive cell ratio compared to control and TGF-β treatment without stimulating cell proliferation (Supplement fig. 1D, E). The results of histology and flow cytometry confirmed the observation of the inhibitory effects of dECM hydrogels on heart fibrosis.

### Microenvironment stiffness modulates the inhibitory effect of dECM hydrogel on fibroblast activation in explant

To study the role of mechanical signals in the natural microenvironment, fibroblast activation was assessed in ventricle explants with altered crosslinking (Fig. 3A). The elastic modulus of decellularized explants after lysyl oxidase inhibitor, BAPN, and ribose treatments was measured by atomic force microscopy (AFM) [13]. AFM confirmed the altered stiffness of the explant extracellular matrix. As described previously, BAPN decreased the elastic modulus from 64kPa to 27kPa, and ribose increased it to 197kPa (Fig. 3B). The density of fibroblasts was not significantly different across the crosslinking groups [13].

**Figure 3.**
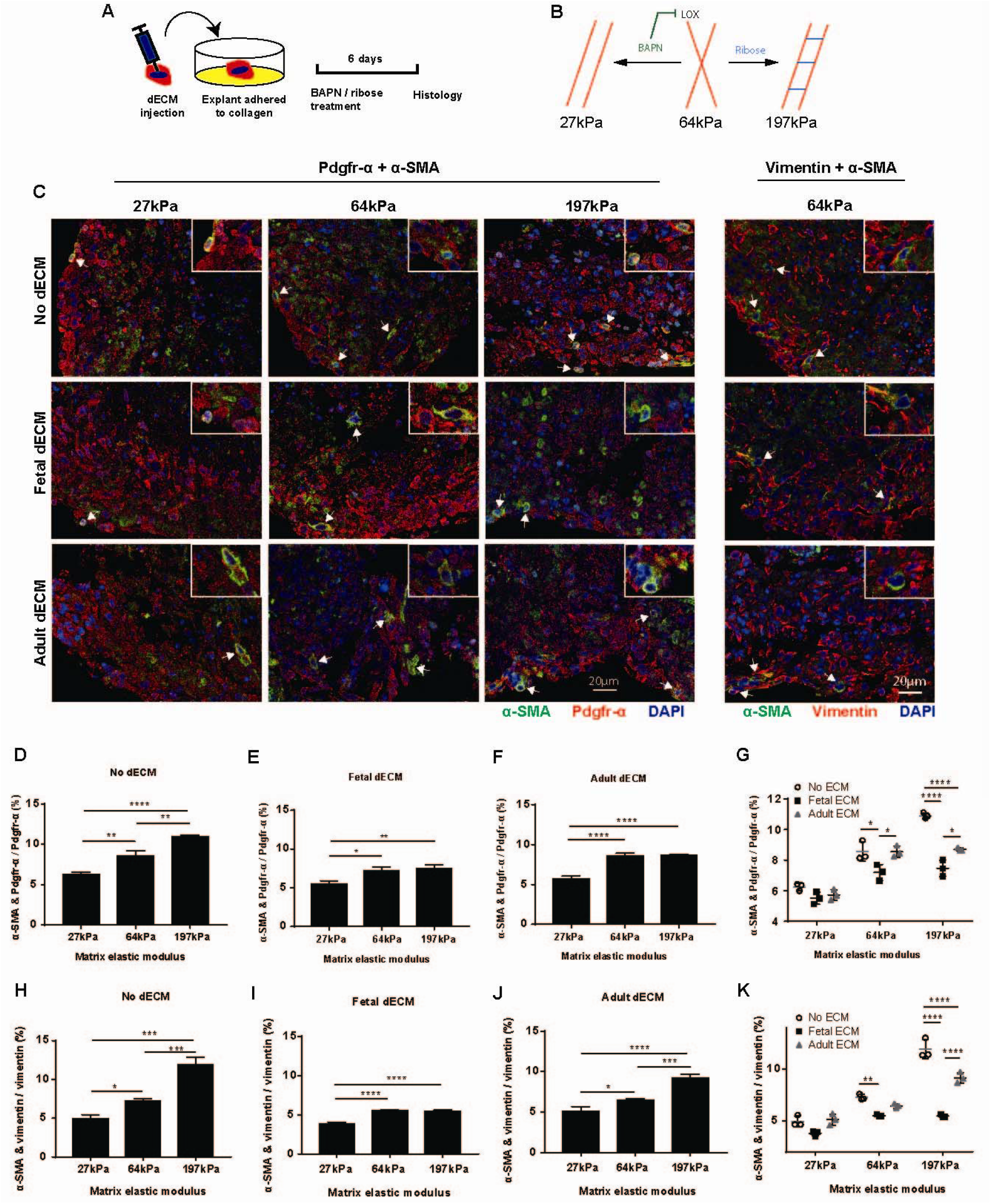
Inhibitory effects of dECM on fibroblast activation in explants are mechanosensitive. (A) dECM hydrogel injected ventricle explants were cultured in media supplemented with BAPN and ribose for 6 days. (B) BAPN lowered decellularized explants stiffness from 64kPa to 27kPa, ribose increased stiffness to 197kPa. (C) Day 1 mouse ventricle explants of different stiffness in combination with adult or fetal dECM hydrogel treatment were fixed and immunostained for α-SMA with vimentin or pdgfr-α. In pdgfr-α positive cells, softening the explants decreased the ratio of α-SMA positive fibroblast in (D) no dECM, (E) fetal dECM, and (F) adult dECM treatments. Stiffening the explants promoted fibroblast activation in no dECM and adult dECM treatments but not in fetal dECM treatment. (G) Adult and fetal dECM treated explants had less α-SMA positive fibroblasts than the control in stiffened 197kPa explant. Fetal dECM, but not adult dECM, lowered α-SMA positive fibroblasts percentage. No significant difference was observed between dECM treated and the control explants in 27kPa softened sample. Similar effects were observed in vimentin positive cells in (H) no ECM, (I) fetal dECM, and (J) adult dECM treatments. (K) Both adult and fetal dECM lowered α-SMA positive fibroblast ratio in stiffened explants but not in the softened ones. Only fetal dECM reduced fibroblast activation in 64kPa explants. (n=3 biological replicates, 3 explants in each biological replicate, one-way ANOVA and Tukey’s test for panel D, E, F, H, I, J, *p<0.05, **p<0.01, ***p<0.001, ****p<0.0001, data presented as mean ± standard deviation. Two-way ANOVA for panel G, K: ECM factor p<0.0001, stiffness factor p<0.0001, interaction p<0.0001, *p<0.05, **p<0.01, ****p<0.0001. White arrow indicates α-SMA and vimentin or pdgfr-α double-positive cell.)

Fibroblast activation was sensitive to the microenvironment stiffness. Fibroblast activation was identified by immunostaining for vimentin or pdgfr-α in combination with α-SMA (Fig. 3C). Softening explants reduced fibroblast activation in the control (Fig. 3D, H), fetal dECM (Fig. 3E, I), and adult dECM (Fig. 3F, J) treated explants. Stiffening explants, in contrast, promoted fibroblast activation in adult dECM treated explants and the control, but not in fetal dECM treated explants. Comparing the fibroblast activation between dECM treated explants and the control demonstrated an inhibitory effect of fetal dECM on fibroblast activation in 64kPa normal and 197kPa stiffened explants (Fig. 3G, K). Fibroblast activation in fetal dECM treated explants was also significantly lower than adult dECM treated samples in the stiffened explants. Adult dECM treatment reduced fibroblast activation in stiffened explants compared to the control but not in the normal and softened explants. The results indicate that dECM treatments limit fibroblast activation in a biologically relevant environment, and the inhibitory effect was more robust in a stiffened microenvironment.

The mechanosensitivity of fibroblasts to dECM treatments was also examined on isolated neonatal rat cardiac fibroblast. Primary cardiac fibroblasts from day 1 rat ventricles were plated on polyacrylamide substrates of various stiffness and treated with adult and fetal dECM (Supplement fig. 2A). Polyacrylamide formulations were confirmed by rheometry (Supplement fig. 2B). The stiffness of the substrates was chosen to mimic the stiffness of mice hearts at different developmental stages [37], [38]. The elastic moduli of the substrates translated from storage modulus are close to the elastic modulus of fetal, neonatal, adult, and fibrotic mice hearts [39]. Increasing substrate stiffness promoted fibroblast activation. The fibroblasts cultured on the highest 21.5kPa substrates showed a higher percentage of activated fibroblast than 0.3kPa and 2.9kPa substrates (Supplement fig. 2C). The stiffness-induced fibroblast activation was reduced with fetal (Supplement fig. 2D) and adult dECM (Supplement fig. 2E) treatments. Fibroblast activation was significantly lowered by fetal dECM treatment on the stiffest (21.5kPa) substrate compared to the control (Supplement fig. 2F). dECM treatments did not change fibroblast activation on the softer (0.3kPa and 2.9kPa) substrates. The results demonstrate that cardiac fibroblast activation stimulated by microenvironment stiffness can be regulated by fetal dECM at stiffness levels approaching fibrotic tissue.

### CAPG and LPXN expression modulated by the combined effects of dECM microenvironment stiffness

Ontological (GO) analysis of mRNA indicated that dECM treatments influence both cytoskeleton organization and cell-matrix binding in ventricle explants (Fig. 4A). Noticeably, a majority of the genes in the top 10 up-regulated GO molecular function categories were related to cell-cell and cell-matrix interaction, whereas many of the genes in the down-regulated categories were related to cytoskeleton dynamics. From the top 100 up and down-regulated genes in the fetal dECM treated explants, 2 genes that regulate cytoskeleton polymerization and expressed in fibroblasts, CAPG and LPXN, were chosen for further analysis. RAC2 was also included because of the correlation of RAC2, CAPG, and LPXN as revealed by protein-protein interaction analysis (Supplement fig. 3). A lower CAPG, LPXN, and RAC2 gene expression levels (one-way ANOVA, Rac2 p-value = 0.049, Lpxn p-value = 0.040, Capg p-value = 0.038) were observed after dECM treatments compared to no dECM treated samples (Fig. 4B).

**Figure 4.**
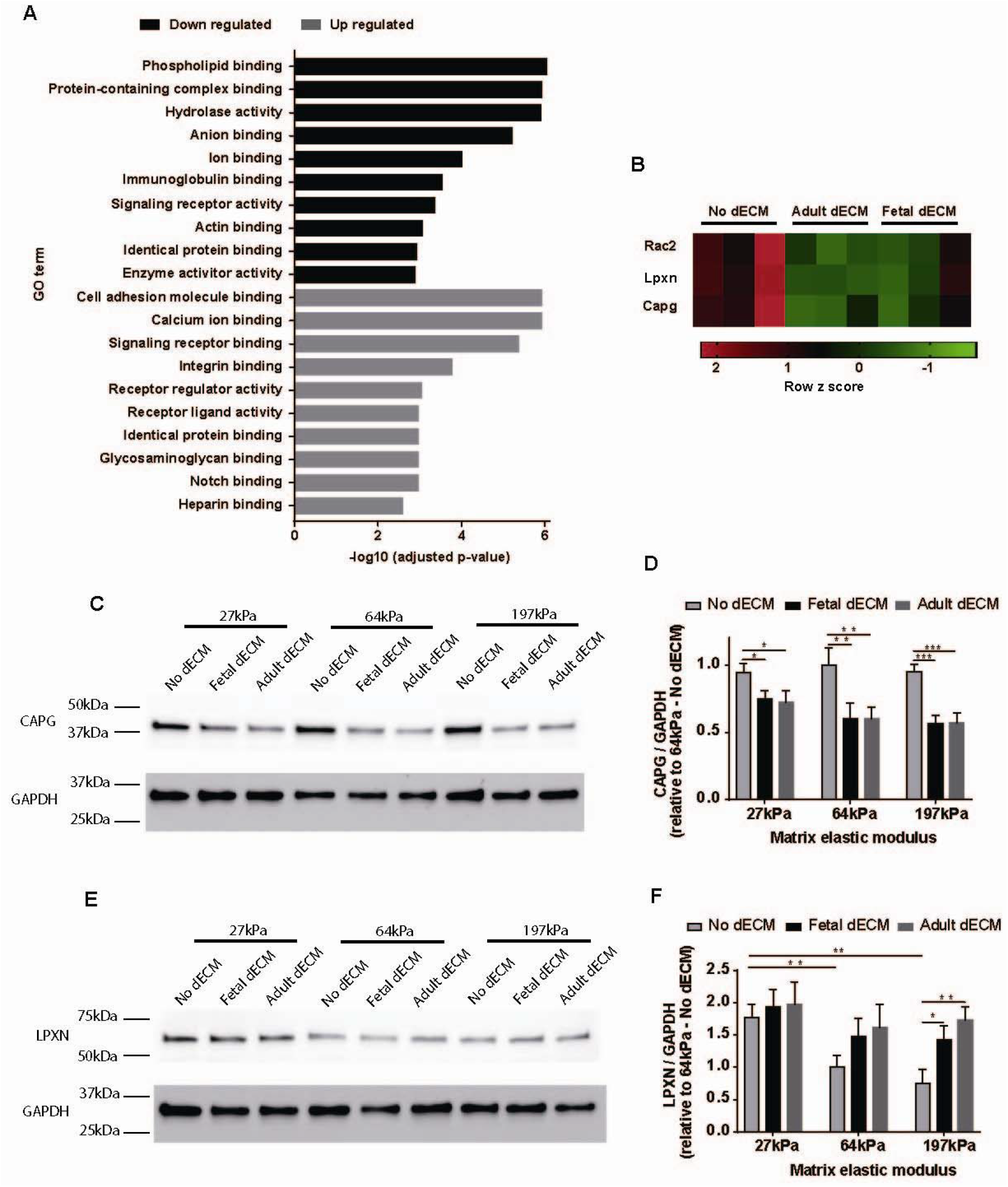
Combined effects of dECM and microenvironment stiffness on CAPG and LPXN expression in explants. (A) GO analysis of differentially expressed genes between fetal dECM treated and the control explants indicated that most of the genes were related to binding. Cytoskeleton organization-related genes were down-regulated, whereas cell-matrix binding proteins were upregulated. (B) CAPG, LPXN, and Rac2 genes were differentially expressed in adult and fetal dECM treated samples. (C) CAPG expression in explants examined by western blot. (D) Both fetal and adult dECM treated explants showed decreased CAPG expression compared to no dECM treated samples. (E) Western blot of LPXN expression demonstrated (F) a lower level in normal (64kPa) and stiffened (197kPa) explants compared to softened explants (27kPa). dECM treatments increased LPXN level only in 197kpa explants. (n=3 biological replicates, 3 explants in each biological replicate, two-way ANOVA and Tukey’s test, panel D: ECM factor p<0.0001, stiffness factor p<0.05, interaction p>0.05. panel F: ECM factor p<0.001, stiffness factor p<0.001, interaction p>0.05. For all panels, *p<0.05, **p<0.01, ***p<0.001, data presented as mean ± standard deviation.)

Western blot analysis of explants indicated an inhibitory effect of dECM hydrogel on CAPG expression (Fig. 4C). CAPG protein level was lowered by both adult and fetal dECM treatments in comparison to no dECM treatment at all tissue stiffness levels (Fig. 4D). LPXN expression was also lowered by increasing stiffness in no-dECM treated explants. Adult and fetal dECM treatments did not significantly change LPXN levels in 27kPa and 64kPa explants. However, LPXN expression was significantly increased by dECM treatments in 197kPa explants (Fig. 4E, F). The western blot results suggest that CAPG expression can be modulated by adult and fetal dECM proteins but not by microenvironment stiffness. In contrast, LPXN expression is sensitive to microenvironment stiffness but only responds to dECM proteins at high stiffness.

### Modulating cytoskeleton affects the inhibitory effect of fetal dECM on fibroblast activation

In order to investigate the function of cytoskeleton in fetal dECM regulated fibroblast activation, explants were treated with actin polymerization inhibitor Latrunculin A (0.1μM), actin polymerization promotor Jasplakinolide (0.1μM), ROCK inhibitor Y27632 (10μM), and focal adhesion kinase (FAK) inhibitor PF573227 (0.1μM) for 3 days (Fig. 5A) and probed with two markers for fibroblasts. No difference was observed in cell morphology and density.

**Figure 5.**
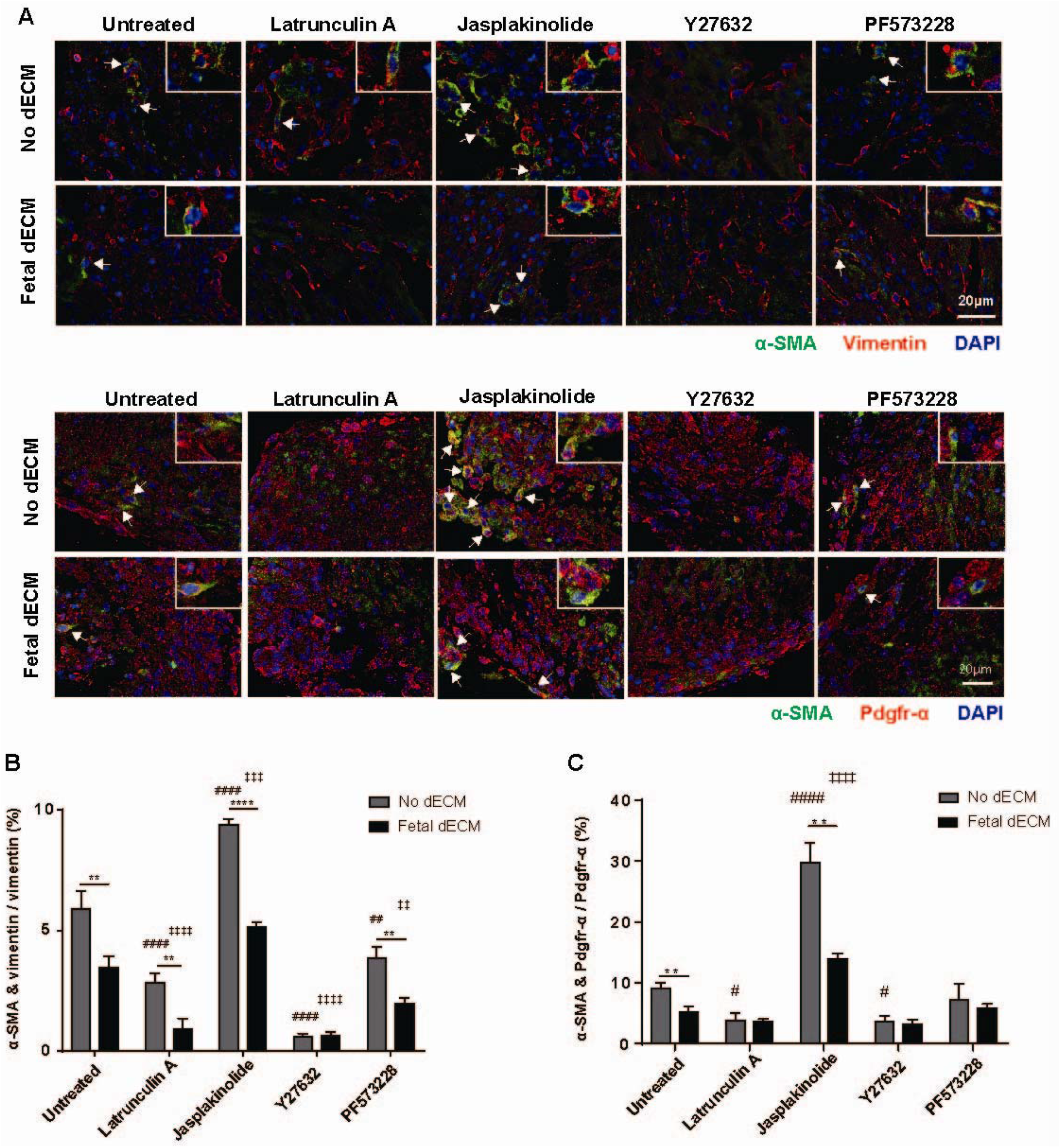
Cytoskeleton mediates the inhibitory effects of fetal dECM hydrogel on fibroblast activation in explants. (A) Day 1 explants were treated by cytoskeleton polymerization modulators and immunostained for α-SMA and vimentin or pdgfr-α. (B) Inhibiting cytoskeleton polymerization with Latrunculin A decreased fibroblast activation in both dECM treated and the control explants. Fetal dECM treatment further lowered the ratio of activated fibroblast compared to the control. A similar effect was observed in PF573228 treated explants. Y27632 treated explants had the lowest ratio of activated fibroblast. Fibroblast activation was not sensitive to fetal dECM in Y27632 treated samples. Jasplakinolide promoted fibroblast activation, but fetal dECM still showed inhibitory effects. (C) In pdgfr-α positive cells, Jasplakinolide stimulated fibroblast activation. Latrunculin-A and Y27632 inhibited fibroblast activation. PF573228, however, did not significantly change fibroblast activation. (n=3 biological replicates, 3 explants in each biological replicate, one-way ANOVA and Tukey’s test for intergroup comparison [e.g. groups in no-dECM treated samples], #p<0.05, ##p<0.01, ####p<0.0001 compared to Untreated no-dECM; ‡‡p<0.01, ‡‡‡p<0.001, ‡‡‡‡p<0.0001 compared to Untreated-Fetal dECM. T-test for intragroup comparison, **p<0.01, ****p<0.0001. Data presented as mean ± standard deviation. White arrow indicates α-SMA and vimentin or pdgfr-α double-positive cell.)

Inhibiting actin polymerization, ROCK inhibition, and focal adhesion perturbation each significantly decreased the activated fibroblast to fibroblast ratio (Fig. 5B, C). In contrast, increasing actin polymerization using Jasplakinolide promoted the percentage of activated fibroblast compared to the untreated group. Fetal dECM treatment significantly lowered fibroblast activation in no dECM and Jasplakinolide treated explants. Fetal dECM also lowered fibroblast activation of vimentin labeled cells after chemical disruption of focal adhesions in explants. Among all the chemical treatment groups, ROCK inhibitor resulted in the lowest activated fibroblast ratio. Fetal dECM did not further reduce activated fibroblasts relative to Y27632 alone (Y27632 no-dECM group). The results demonstrate that the inhibitory effect of fetal dECM on fibroblast activation is sensitive to actin integrity and focal adhesion activity.

Ventricle explants were then probed for CAPG and LPXN to determine if these two proteins were expressed in cardiac fibroblast and for evidence of a role in dECM hydrogel reduction of fibroblast activation (Fig. 6A). CAPG protein is primarily expressed in fibroblasts in the heart. Disrupting ROCK, FAK, and cytoskeleton polymerization lowered the percentage of CAPG positive fibroblast in explants (no dECM). Chemically stabilizing cytoskeleton polymerization induced the reverse effect of increasing CAPG positive cells. Fetal dECM significantly reduced CAPG positive cell ratio compared to no dECM treated samples only in untreated and Jasplakinolide groups which have normal and enhanced actin polymerization, respectively (Fig. 6B). LPXN is expressed in various cell types including fibroblast. Fetal dECM increased LPXN positive fibroblast ratio compared to no dECM treated samples in all chemical treatment groups except Y27632. Inhibiting FAK activity increased the percentage of LPXN positive fibroblast in comparison to untreated explants with or without dECM treatment (Fig. 6C). The results suggest that CAPG and LPXN are effectors of exogenous dECM hydrogel in cardiac fibroblast and CAPG level is positively related to fibroblast activation.

**Figure 6.**
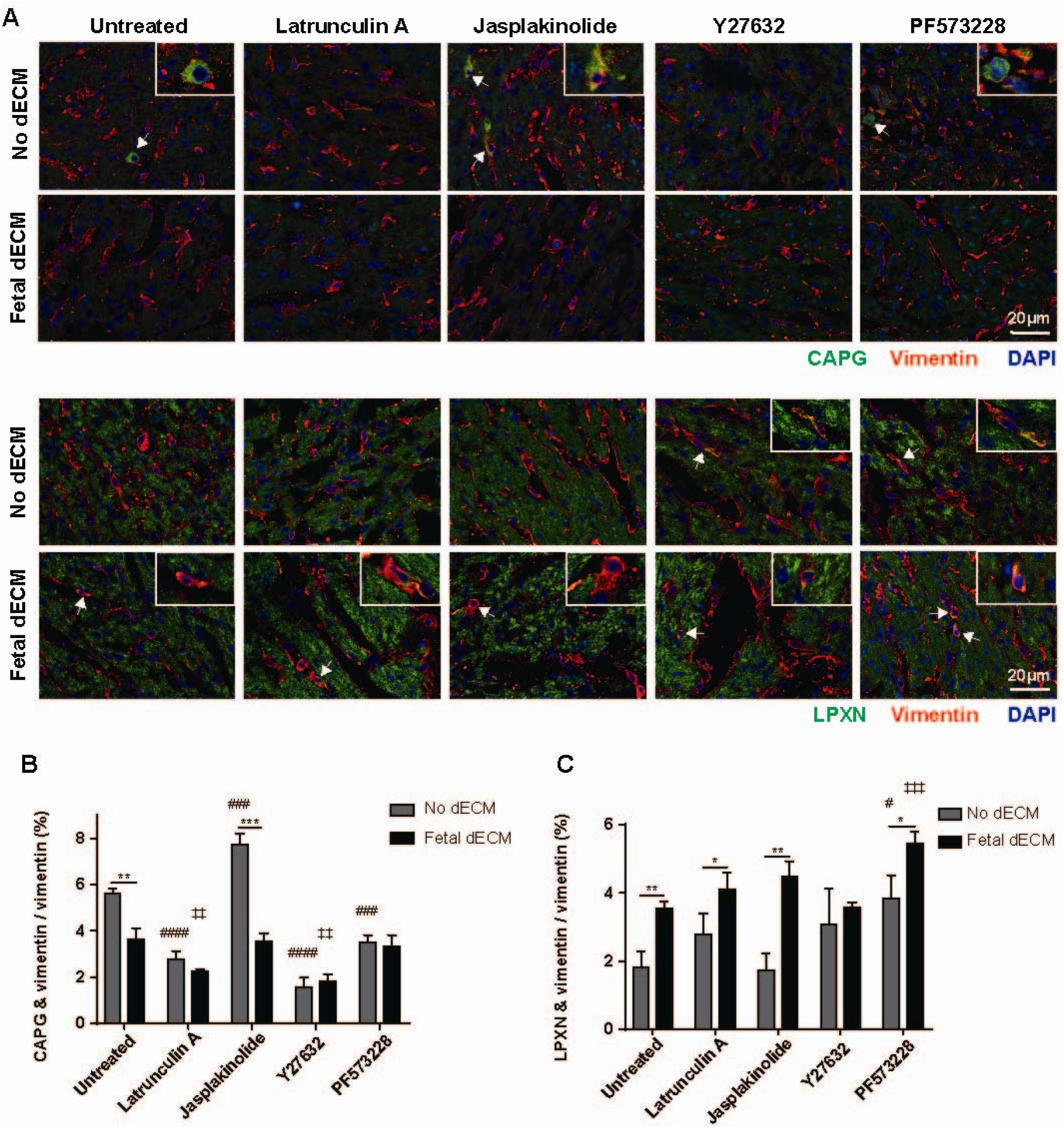
Modulating cytoskeleton polymerization and FAK changes CAPG and LPXN expression in explants. (A) CAPG and LPXN localization in ventricle explants were examined by immunostaining. (B) Inhibiting cytoskeleton polymerization and FAK activity decreased the ratio of CAPG positive fibroblasts in explants. Facilitating cytoskeleton polymerization showed a reverse effect. Fetal dECM decreased CAPG positive fibroblast ratio in untreated and Jasplakinolide treated samples compared to no dECM treatment. (C) LPXN positive fibroblast ratio was increased by FAK inhibition but not affected by cytoskeleton polymerization in no dECM treated samples. Fetal dECM treatment increased LPXN positive cell number compared to no dECM treated explants in all chemical groups except Y27632. (n=3 biological replicates, 3 explants in each biological replicate, one-way ANOVA and Tukey’s test for intergroup comparison [e.g. groups in no-dECM treated samples], #p<0.05, ###p<0.001, ####p<0.0001 compared to Untreated-No dECM; ‡‡p<0.01, ‡‡‡p<0.001 compared to Untreated-Fetal dECM. T-test for intragroup comparison, *p<0.05, **p<0.01. Data presented as mean ± standard deviation. White arrow indicates CAPG/ LPXN and vimentin double-positive cell.)

The influences of cytoskeleton modulators on α-SMA, CAPG, and LPXN expression in no-ECM treated fibroblasts were examined in primary fibroblasts isolated from day 1 neonatal rat hearts (Supplement fig. 4A). Similar to fibroblasts in ventricle explants (by histology), primary fibroblasts cultured in dish showed increased fibroblast activation after jasplakinolide treatment and lowered ratio after actin destabilization (latrunculin A) and ROCK inhibiton (Y27632) treatments (Supplement fig. 4B). Perturbing focal adhesion activity (PF573228 treatment) did not significantly change fibroblast activation. CAPG positive cell percentage was reduced by latrunculin A, Y27632, and PF573228, and increased by jasplakinolide (Supplement fig. 4C). A higher LPXN positive cell ratio was observed in PF573228 treated cells (Supplement fig. 4D). The other chemicals did not change LPXN positive cell number compared to the control. The cell culture experiments indicate that cytoskeleton modulators induce similar changes in fibroblast cultured in dish and in explants for α-SMA, CAPG, and LPXN expression.

### Decreasing CAPG expression blocks activation of cardiac fibroblast

CAPG expression was lowered by CAPG siRNA to determine if CAPG regulates cardiac fibroblast activation. Treating cardiac fibroblasts with CAPG siRNA decreased CAPG protein level by 60.3% (Fig. 7A, B). Interfering CAPG expression significantly decreased the ratio of α-SMA positive fibroblasts compared to no siRNA treatment in the control and TGF-β treated cells (Fig. 7C, D). Fetal dECM treated fibroblasts showed a lower ratio of α-SMA positive fibroblasts than the untreated control. Interfering CAPG expression in fetal dECM treated cells did not further limit fibroblast activation which suggests CAPG activity had been lowered by dECM and thus further decreasing CAPG level by siRNA did not affect fibroblast activation. To determine if CAPG knock-down also affects collagen production, collagen type Iα1 expression in primary fibroblasts was examined by western blot (Fig. 7E). CAPG siRNA treatment lowered collagen Iα1 level compared to control (Fig. 7F). The results indicate dECM hydrogel lowered fibroblast activation through CAPG.

**Figure 7.**
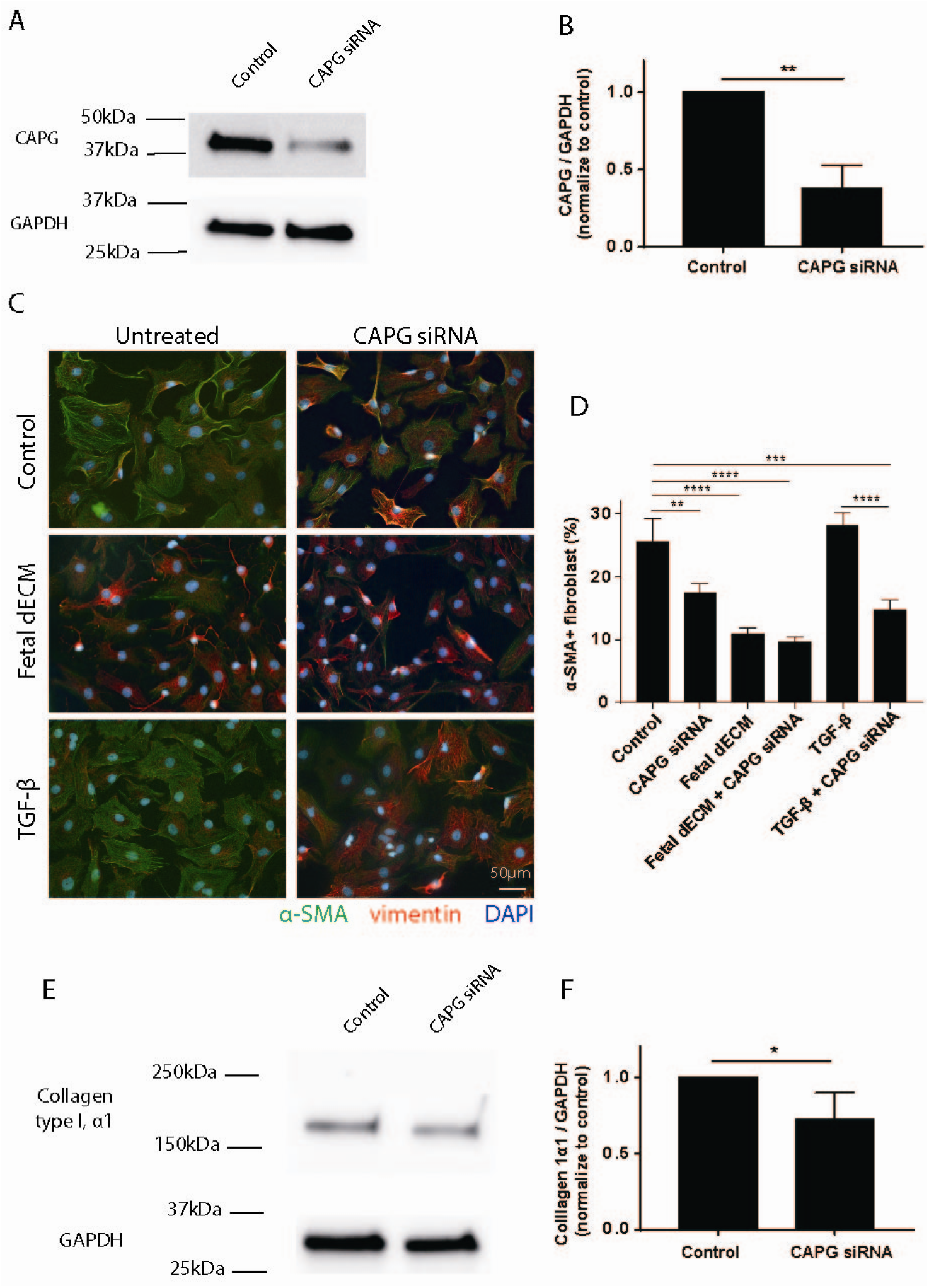
Knocking down CAPG expression inhibits fibroblast activation in dish. (A) Fibroblast CAPG expression was evaluated by western blot after 48h CAPG siRNA treatment. (B) CAPG expression was significantly decreased by siRNA treatment. (C) Fibroblast activation after CAPG knockdown was examined by immunostaining. (D) Decreasing CAPG expression significantly lowered the ratio of α-SMA positive fibroblast in TGF-β treated cells and control. (E) Collagen type Iα1 expression was examined by western blot. (F) Knock down of CAPG level lowered collagen 1α1 expression in fibroblasts compared to control. (panel B, F: n=3 biological replicates, 2 technical replicates in each biological replicate, t-test, *p<0.05; panel D: n=3 biological replicates, 4 technical replicates in each biological replicate, one-way ANOVA and Tukey’s test, *p<0.05, **p<0.01, ***p<0.001, ****p<0.0001. Data presented as mean ± standard deviation.)

## Discussion

This work demonstrates that exogenous decellularized extracellular matrix (dECM) limits collagen deposition and fibroblast activation in cardiac tissue through actin regulatory protein CAPG. Reducing extracellular stiffness enhanced the dECM anti-fibrotic effects. Previous studies have shown that increasing substrate stiffness enhances TGF-β1 induced fibroblast activation [10], that matrix stiffness affects fibroblast activation induced by cancer cell generated microvesicles [40], that stiffness alone regulates fibroblast pro-fibrotic phenotype, migration, and function [8], [41]–[43], and that fibroblast activation can be significantly reduced by Rho-kinase inhibition [44]–[46]. Nevertheless, reducing cardiac fibroblast activation and collagen deposition using naturally derived biomaterial has not been reported. We are the first to indicate that cardiac extracellular matrix derived hydrogel inhibits fibrosis in cardiac tissue. Further, CAPG has not been reported as a target for fibroblast activation inhibition. While the expression of dECM-mediated CAPG appears to lack mechanosensitivity, focal adhesion associated LPXN expression was altered by changes in stiffness. This study suggests that cardiac dECM hydrogel may be used as a therapy for heart ischemic injury and fibrosis.

Extracellular matrix derived biomaterials have been demonstrated to improve cardiac function post-heart injury in animal models. The mechanism of dECM induced heart regeneration has not been fully understood. We show that the dECM effects include reduced fibroblast activation and collagen deposition in post-MI hearts and in heart explants. Thus heart explants can be used as an *in vitro* model of heart post-injury response that retains the natural 3D-environment of heart and can be cultured in controlled conditions. Investigating dECM treated explants may reveal the signaling pathways that regulate heart regeneration. Multiple extracellular matrix proteins, like neuregulin, agrin, periostin, and follistatin-like 1, have been reported to promote cardiomyocyte proliferation or angiogenesis in the injured adult mice hearts [47]–[51]. The extracellular matrix ligands that reduce cardiac fibrosis have not been well identified. TGF-β, angiotensin II, connective tissue growth factor, and platelet-derived growth factor have been examined as targets for fibrosis inhibition [52]–[57]. However, because most of the fibrosis regulators are involved in multiple biological processes, inhibiting the factors may not significantly improve cardiac function or cause severe side effects such as increased mortality [58]. Our results suggest that exogenous fetal cardiac dECM hydrogel attenuates fibroblast activation and cardiac fibrosis without observable side effects. In order to determine if dECM hydrogel can be developed to treatment for heart fibrosis, *in vivo* experiments in heart-injury model are needed. Unlike fetal dECM, adult dECM lowered fibroblasts activation only in stiffened heart explants. This is possibly due to the different components in adult and fetal dECM. The adult and fetal extracellular molecular profiles of the heart have distinct protein composition despite significant overlap of the proteome [59]. The ratio of cardiac matrix proteins, such as collagen, fibronectin, and agrin, changes significantly from young developmental age to adulthood [13], [59]. We previously confirmed higher agrin levels in fetal dECM among other proteins can drive signaling regenerative signaling in cardiomyocytes [13]. Further characterization of the extracellular proteome will identify specific molecules that mediate differential responses of adult and fetal dECM on fibroblast activation.

Stable tissue stiffening is generally preceded by reactive fibrosis in adult post-MI hearts unlike the transient reversible response in early-age neonates. It has been reported that myocardial stiffness is primarily determined by fibrosis, and not by cardiomyocyte hypertrophy in hypertensive heart [60]. The infarct area in heart, which is composed mainly of fibrotic tissue, has 3 to 6 fold increase in stiffness compared to healthy heart tissue [61]–[64]. It is not clear if increased myocardial stiffness directly impairs the cardiac regenerative capacity in adult mammals. Increasing matrix stiffness promotes fibroblast activation and proliferation [41]. Conversely, decreasing heart stiffness reduces fibrosis [29]. Lowering microenvironment stiffness also promotes cardiomyocyte division and fetal dECM-induced cardiomyocyte cell cycle activity [13], [65]. The use of BAPN as a softening agent is a confounding factor in explants since it inherently interrupts tissue remodeling. Alternative approaches to mechanically unloading the heart or softening the cell niche will strengthen understanding of the observed phenomena in animal models. We previously showed that primary cardiomyocytes on elastomeric substrates exhibit mechano-sensitivity to the dECM treatments. We also observed by histological analysis that fetal dECM reduced collagen deposition in stiffened heart explants [13]. Here, we investigate the distinctive phenomena of matrix modulation of stiffness-activated fibroblasts at cellular resolution. We observe fetal dECM and softening contribute to reducing basal fibroblast activation in ventricle explants. In contrast, adult dECM shows inhibitory effects to activation only in stiffened explants. It is not clear if the dECM inhibitory effects on activation of fibroblast occurs through direct action of soluble factors or insoluble components of the dECM. Further investigations are required to determine the influences of extracellular interactions on fibroblast activation with dECM treatment. Of consequence, the fibroblast activation and transcriptome analysis were examined after formation of a robust fibrotic area. Investigating the fibroblasts and remodeling dynamics at earlier time points will reveal more information about the synergistic effect of dECM and tissue stiffness on fibroblast activation and fibrotic protein deposition.

In this study we investigated whether physical changes associated with age and disease can alter physiological responses to extracellular proteins that modulate fibroblast activation to a fibrotic state. Our data suggest cytoskeletal integrity can modulate cellular responses to fetal ‘regenerative’ signals. Here, we hypothesize that extracellular driven signaling intersects with mechanotransduction signaling pathways through cytoskeletal signaling in CAPG and focal adhesion in LPXN in fibroblasts. Rho-kinases have been reported to mediate fibrosis in multiple organs [66]–[69]. Rho-associated kinases regulate fibroblast contractile force generation, migration, activation, and proliferation [45], [70]–[72]. Our results indicated that dECM down-regulates a rho family kinase, rac2 related signaling pathways, specifically CAPG and LPXN. LPXN is homologous to a focal adhesion protein, paxillin. Paxillin-null fibroblasts show some defects in cell spreading and altered cell motility in response to physical stimuli [73], [74]. The uncorrelated LPXN mRNA and protein levels can be caused by post-translation modifications [75], [76]. Nevertheless, both data suggest LPXN expression is changed by dECM treatments and changes in stiffness. Phosphorylation and translocation also affects LPXN activity [23],[77]. Investigating LPXN activity will determine its role in regulating fibroblast activation and possible interaction with CAPG signaling. CAPG is a member of gelsolin/villin family of actin regulatory proteins. Gelsolin depletion inhibits force-induced α-SMA promoter activation and cell migration in fibroblasts [78], [79]. It has been reported to control ruffling in different cell types. CAPG null engineered mice are viable without robust morphological deformities; however, there is altered macrophage activity [80]. Our results demonstrate that CAPG expression is positively correlated to cardiac fibroblast activation. We show that both fetal dECM and independently CAPG depletion with siRNA decrease stiffness-mediated fibroblast activation in a manner that resembles pharmacological disruption of cytoskeleton both in explants and primary cells. Similarly, we observe that pharmacological stabilization of the cytoskeleton both enhances fibroblast activation and increases CAPG expression. CAPG depletion also reduces collagen expression. It remains to be determined if gain and loss of CAPG expression in cardiac fibroblasts can alter fetal dECM effects on the post-ischemic response. Thus, CAPG may be developed as a target for anti-fibrosis treatment in the diseased heart and has shown promise as a target for cancer [18].

This study provides some insight into molecular mechanisms of dECM treatment and microenvironment stiffness regulation of post-MI fibrosis. A noted confounding factor is the heterogeneity of fibroblasts, identified only by vimentin and pdgfr-α markers, and identification of activated fibroblasts solely by α-SMA and contractility. Alpha smooth muscle actin is a common marker of fibroblast activation demonstrating reorganization of the cytoskeleton; however it does not always correlate with the myofibroblast proliferation and matrix secreting phenotype. While we do observe reduced collagen I expression with CAPG knock down, broader analysis of matrix secretion of collagen subtypes and periostin, and extracellular enzymes such as matrix metalloproteinases would be helpful in our interpretation. As noted, proteomic analysis of the dECM treatments would also help in mapping the activators of the two genes analyzed here. The correlation between CAPG expression and fibroblast activation *in vivo* may yield new information on dynamics of fibroblast activation. Further investigations are required to understand the effects of CAPG and LPXN knock-down and overexpression in animal models and the role in cardiac fibroblasts.

## Conclusion

This study confirms that increased microenvironment stiffness on 2D substrate and in 3D ventricle explants promotes fibroblast activation. Fetal cardiac dECM hydrogel inhibits fibroblast activation at all stiffness levels in explants. Adult dECM hydrogel lowers fibroblast activation only in stiff explants. dECM hydrogel inhibits fibroblast activation through Rho-kinases and CAPG protein.

## Acknowledgment

This work was supported by Faculty Investment Fund RES221997 from Case Western Reserve University (CWRU) (S.E.S), NIH 1 C06 RR12463-01, and R01EY021731 (P.S-.H.P). We would like to acknowledge the use of microscopes in the Light Microscopy Imaging Facility at CWRU, made available through NIH Grant S10-OD024996. We thank Akinola Akinbote and Bhargavee Gnanasambandam for preliminary experiments with primary fibroblasts. We thank Gabrielle McBroom for mRNA data analysis. We thank Dr. Nicholas P. Ziats and Sandra Siedlak for assistance with histology. We thank Craig Watson, Ali Ansari, and Chao Liu for providing technical support. We thank Dr. Timothy Meade and Dr. Suneel Apte for technical support with cell contractility.

## Data availability statement

The datasets generated during the current study are available from the corresponding author by reasonable request.

