## Supplemental documents for "Exogenous extracellular matrix proteins decrease cardiac fibroblast activation in stiffening microenvironment through CAPG"

**Supplementary material**

**Methods**

**Rheometry.** The storage modulus of polyacrylamide gels was measured using MARS III rheometer (HAAKE). A 35mm plate was used for plate/plate measurement. Storage modulus was probed using frequency sweep from 100Hz to 0.1Hz at 1% strain at 37°C.

**Gel contraction assay.** A mold made of agarose (3%; MIDSCI) was sterilized and added to a 24-well plate where a cloning cylinder was placed to make a 10 cm hole. The mold was equilibrated in DMEM. One-part NIH 3T3 cells ($3\times{10}^{5}$ cells/mL) in 10% FBS DMEM (5mM glucose) was mixed with one-part neutralized 2.4mg/ml collagen mixture to a final concentration of 1.2 mg/mL of collagen and $1.5\times{10}^{5}$ cells/mL. The solution was added to mold and incubated for 1 hour at 37°C. After matrix polymerization, 10% FBS DMEM was added to each group: low glucose DMEM (5mM), low glucose + TGF-β (10ng/mL; PeproTech), low glucose + fetal dECM (200μg/ml). Then, the collagen disc was dislodged from the inner surface of the agarose mold and incubated in 37°C. Images were taken every 24h, starting from 0h, for 4 days. During the incubation period, the media was not changed, and no additional dosage of treatment occurred. ImageJ software (NIH) was used to measure the total are of the gel. The scale was set from a 10 mm reference line drawn with a ruler. Each gel area was normalized to its 0-hour measurement.

**Flow cytometry.** NIH 3T3 cells were plated in cell culture dish with a seeding density of 7,000 cells per cm^2^ and cultured in 5% FBS DMEM for 30 hours. After washing once with 1X PBS, cells were cultured with different conditions: 1% FBS DMEM (Control group), 10 ng/mL TGF-β (PeproTech 10021) in 1% FBS DMEM (TGF-β group), or 100 µg/mL fetal dECM in 1% FBS DMEM (Fetal dECM group) for 24 hours. Cells were dissociated with 0.25% Trypsin-EDTA (Gibco). Cells were fixed in 4% paraformaldehyde for 15 minutes and permeabilized using permeabilization buffer (0.3% Triton X-100 and 0.5% Bovine Serum Albumin in 1X PBS) for 10 minutes. Cells were immunostained with α-SMA vimentin primary antibodies 1:200 diluted in antibody dilution buffer (0.5% Bovine Serum Albumin in 1X PBS) for 1 hour at RT, and incubated in Alexa Fluor conjugated secondary antibodies for 30 minutes. Immunofluorescent labeled cells were resuspended in 1XPBS with 1 million cells per mL solution, then cells were analyzed by flow cytometry. All sample acquisitions were performed on LSR II flow cytometer (BD Biosciences), and analyses were performed using the FlowJo software. Side-scatter and forward scatter profiles were used to eliminate cell doublets and debris.

**Polyacrylamide substrate preparation.** Polyacrylamide preparation was adapted from previously published work to achieve substrates of varying compliance [32]. In brief, acrylamide, bis-acrylamide, ammonium persulfate, and TEMED were mixed and diluted to the desired concentration and dropped on ozone cleaned coverslip. Polyacrylamide solution was dispensed on round coverslips and allowed to polymerize at room temperature before washing in PBS. Collagen type I was linked to polyacrylamide substrate by sulfo-SANPAH.

**Primary fibroblasts culture.** Fibroblasts isolated from day 1 neonatal rats were passaged and expanded in cell culture flasks twice. Cells were plated to 48-well plates at 20,000 cells per well. After overnight plating in 5% FBS-DMEM media, cells were washed in 1xPBS once and cultured in DMEM supplemented with Latrunculin A (0.1μM), Jasplakinolide (0.01μM), Y27632 (10μM), and PF573228 (0.1μM) for 48h. Cells were then fixed and immunostained.

**Figures**

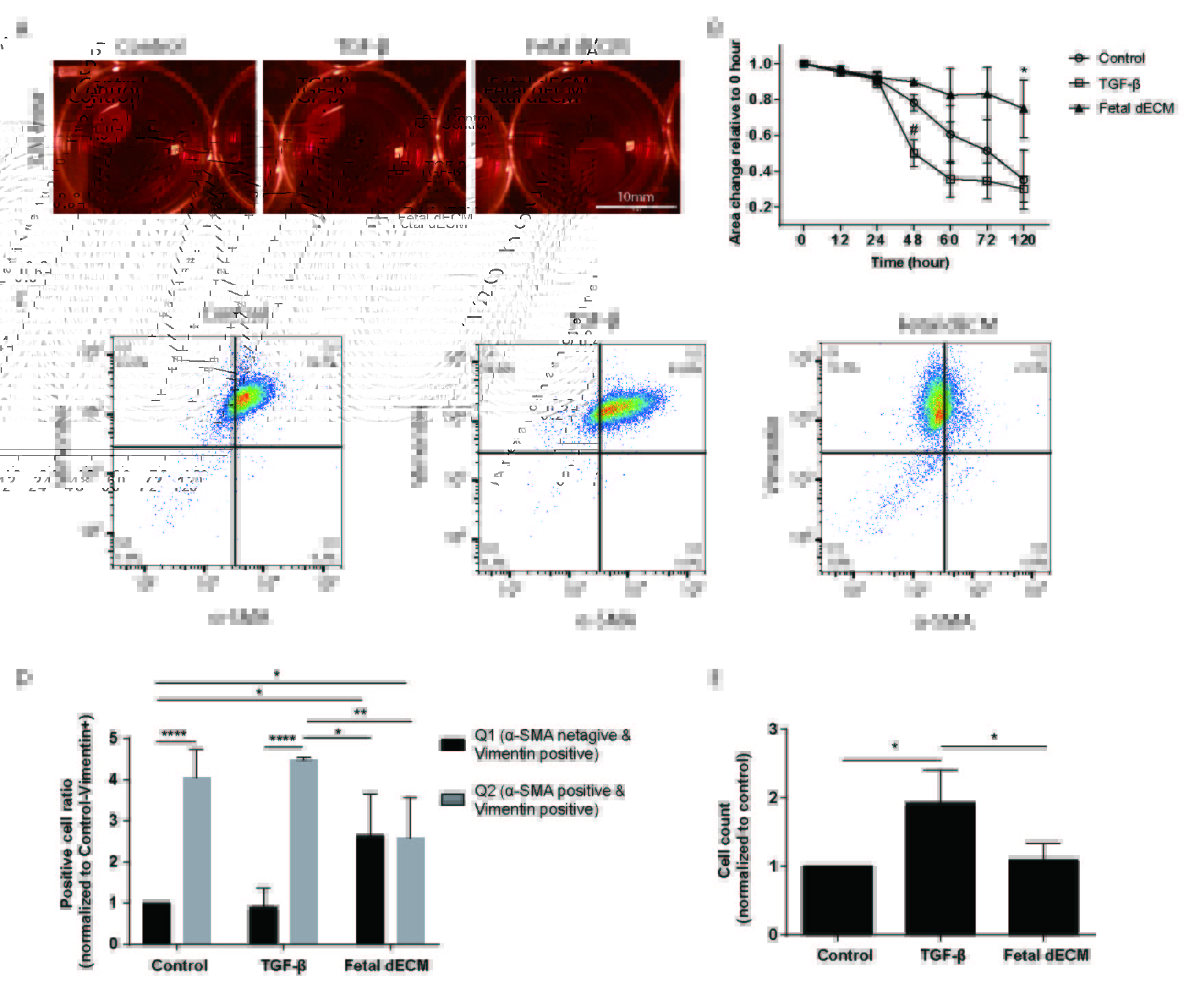

**Supplementary figure 1**. **The dECM treatment inhibits fibroblast contractility and α-SMA expression.** (A) 3T3 cells culture in collagen type I gel for 120 hours with untreated media, fetal dECM and TGF-β. (B) Fetal dECM treatment lowered cell-mediated collagen gel contraction compared to the control. TGF-β treatment (positive control) increased contractility. (C) α-SMA positive cells were counted by flow cytometry. (D) Fetal dECM treatment lowered percentage of α-SMA positive cells compared to control and TGF-β treatment. (E) TGF-β significantly increased the cell number; while dECM lowered basal α-SMA without affecting cell number. (panel B: n=3 biological replicates, 3 technical replicates in each biological replicate, one-way ANOVA and Tukey’s test, *p<0.05 for Fetal dECM vs. Control, #p<0.05 for TGF-β vs. Control. panel D: n=4 biological replicates, two-way ANOVA and Tukey’s test, *p<0.05, **p<0.01, ****p<0.0001. panel E: n=4 biological replicates, one-way ANOVA and Tukey’s test, *p<0.05. Data presented as mean ± standard deviation.)

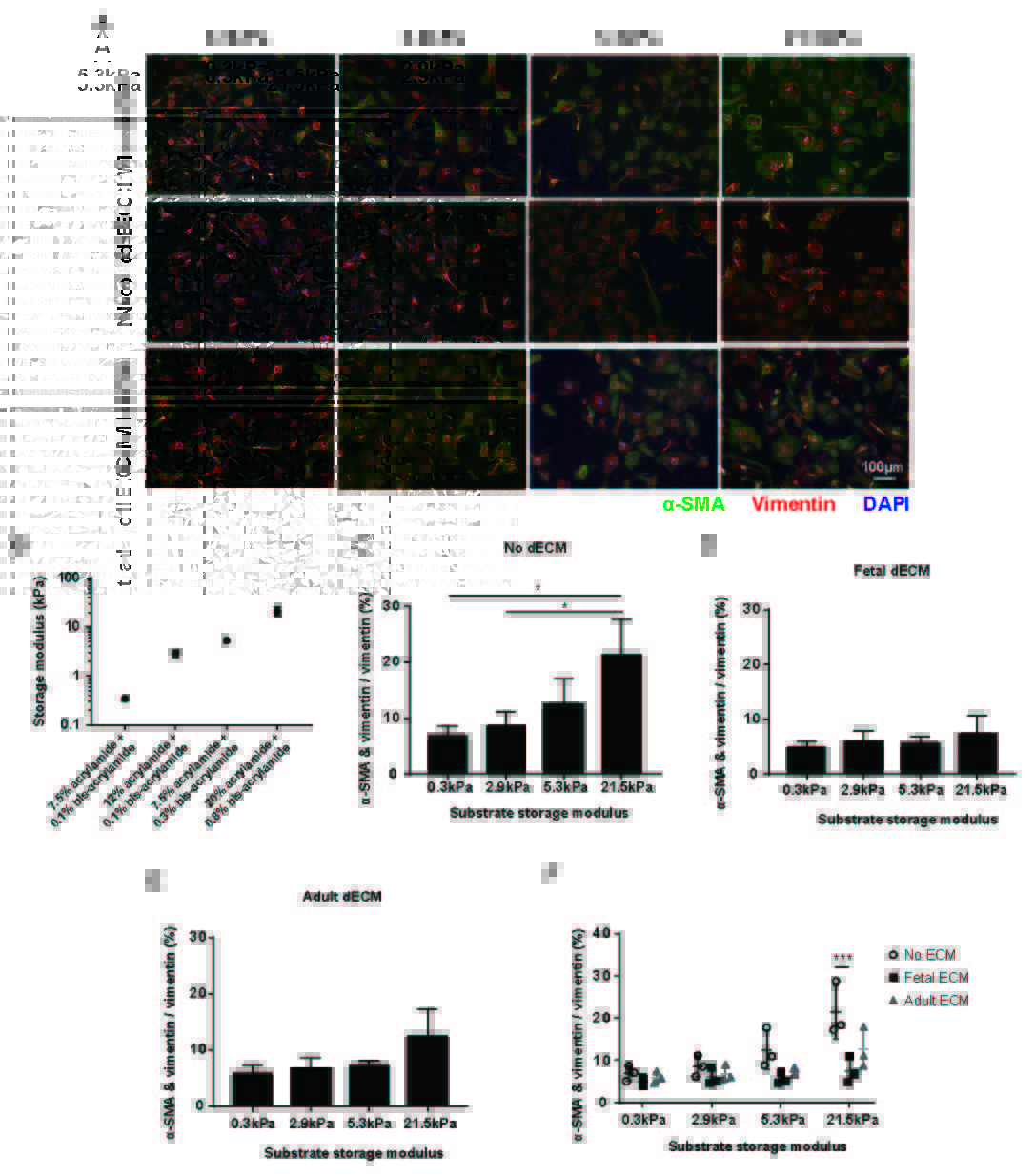

**Supplementary figure 2. dECM treatments reduced stiffness-mediated fibroblast activation on elastomeric substrates.** (A) Fibroblasts cultured on polyacrylamide substrates of varying stiffness were immunostained for α-SMA and vimentin. (B) Storage modulus of polyacrylamide gels at 1Hz sweep rate. (C) Stiffness increased the percentage of α-SMA positive fibroblasts in the no-dECM treated groups. Substrates stiffness effects on α-SMA expression were blocked with (D) fetal dECM and (E) adult dECM treatment. (F) Adult dECM treatment significantly lowered fibroblast activation on the stiffest (21.5kPa) substrate in comparison to the control. Fetal dECM further reduced fibroblast activation compared to adult dECM. (panel B: n=3 replicates at different time points, one sample measured each time. Panel C, D, E: n=3 biological replicates, 3 explants in each biological replicate, one-way ANOVA and Tukey’s test, *p<0.05, **p<0.01, ***p<0.001, ****p<0.0001. Panel F: Two-way ANOVA and Tukey’s test: ECM factor p<0.001, stiffness factor p<0.0001, interaction p≤0.05, ***p<0.001. Data presented as mean ± standard deviation.)

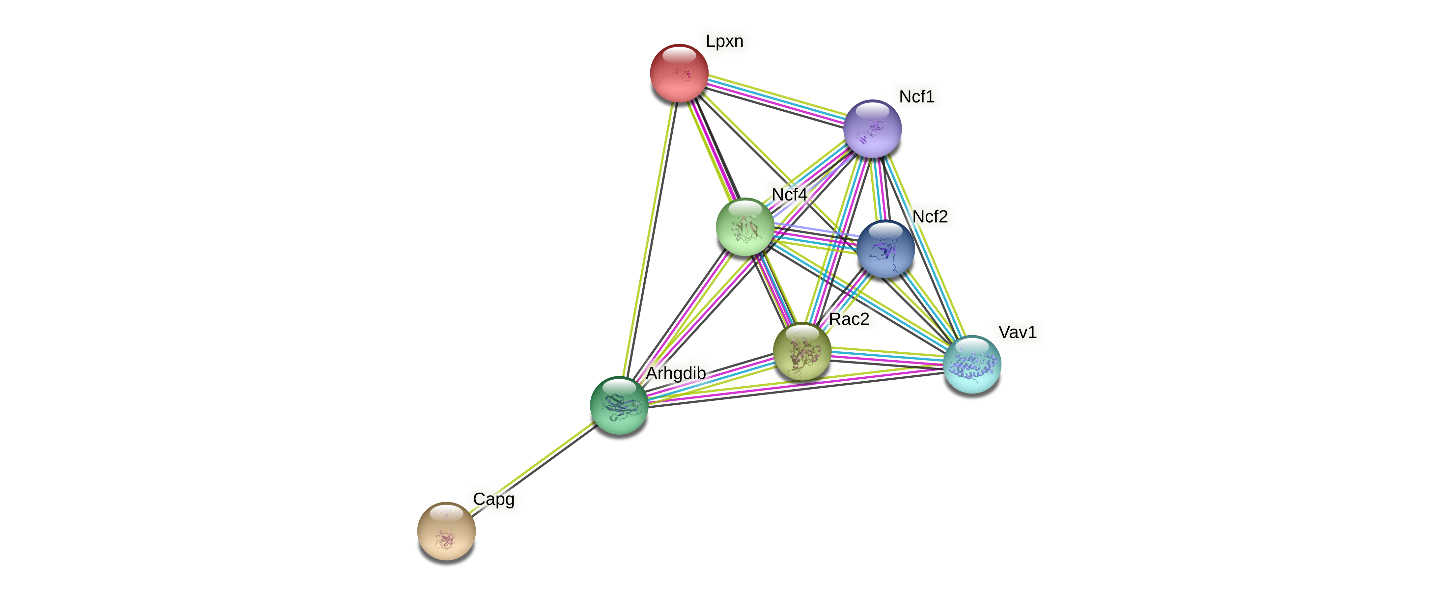

**Supplementary figure 3. Possible signaling pathway connecting CAPG, LPXN, and RAC2 derived from the String v11 protein-protein association network.**

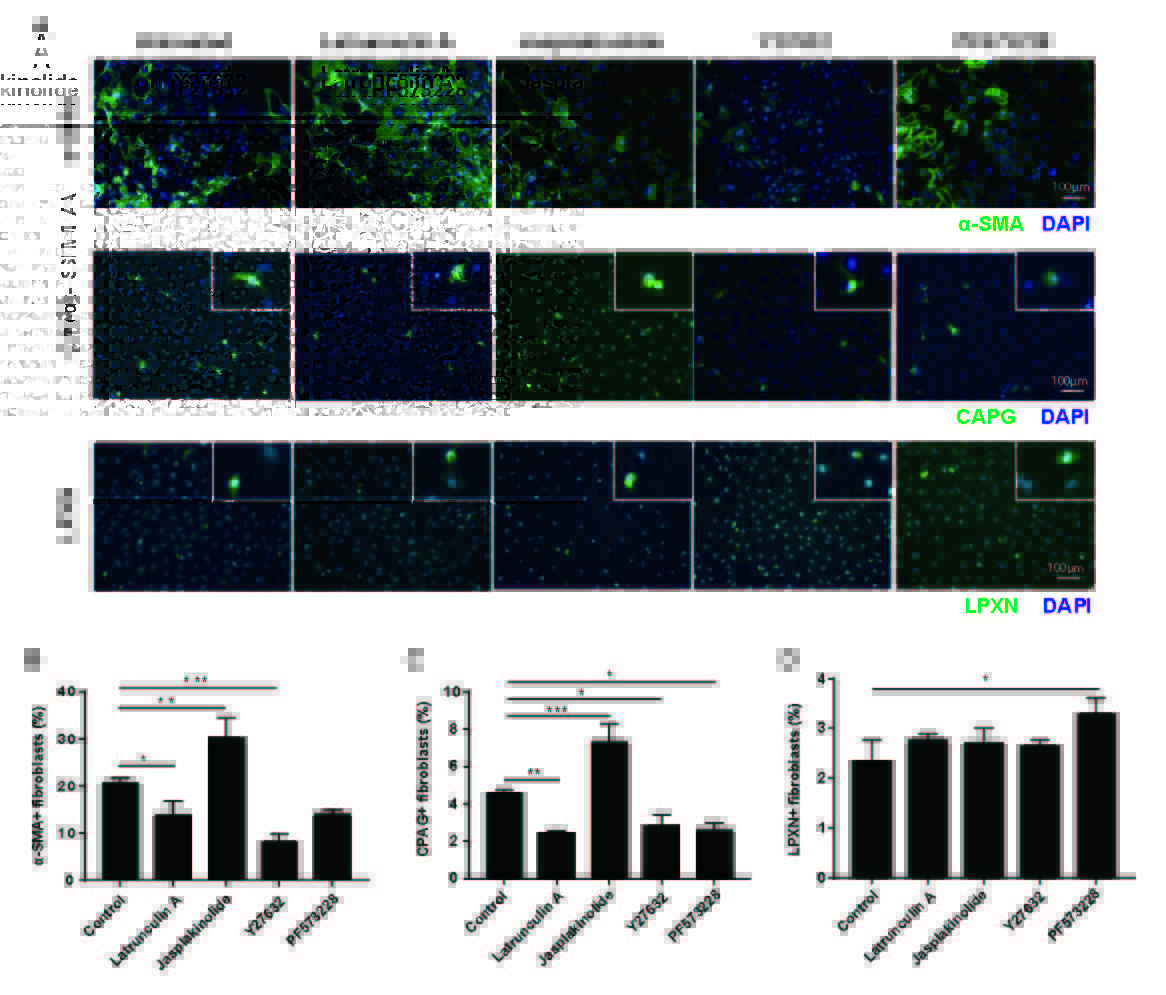

**Supplementary figure 4. Modulating cytoskeleton polymerization in primary fibroblasts changes α-SMA, CAPG, and LPXN expression.** (A) α-SMA, CAPG, and LPXN expression in fibroblasts were examined by immunostaining. (B) Disrupting cytoskeleton polymerization using latrunculin A and inhibiting ROCK activity using Y27632 lowered α-SMA positive cells compared to the control. Increasing cytoskeleton polymerization using Jasplakinolide promoted α-SMA positive cell number. Inhibiting FAK activity using PF573228 did not significantly change α-SMA expression. (C) CAPG positive cell number was lowered by Latrunculin A, Y27632, and PF573228 compared to the control. Jaskplakinolide, on the other hand, increased CAPG positive cell number. (D) LPXN positive cell percentage was increased by PF573228 treatment alone compared to the control. (n=3 biological replicates, 4 technical replicates in each biological replicate, one-way ANOVA and Tukey’s test, *p<0.05, **p<0.01, ***p<0.001. Data presented as mean ± standard deviation.)

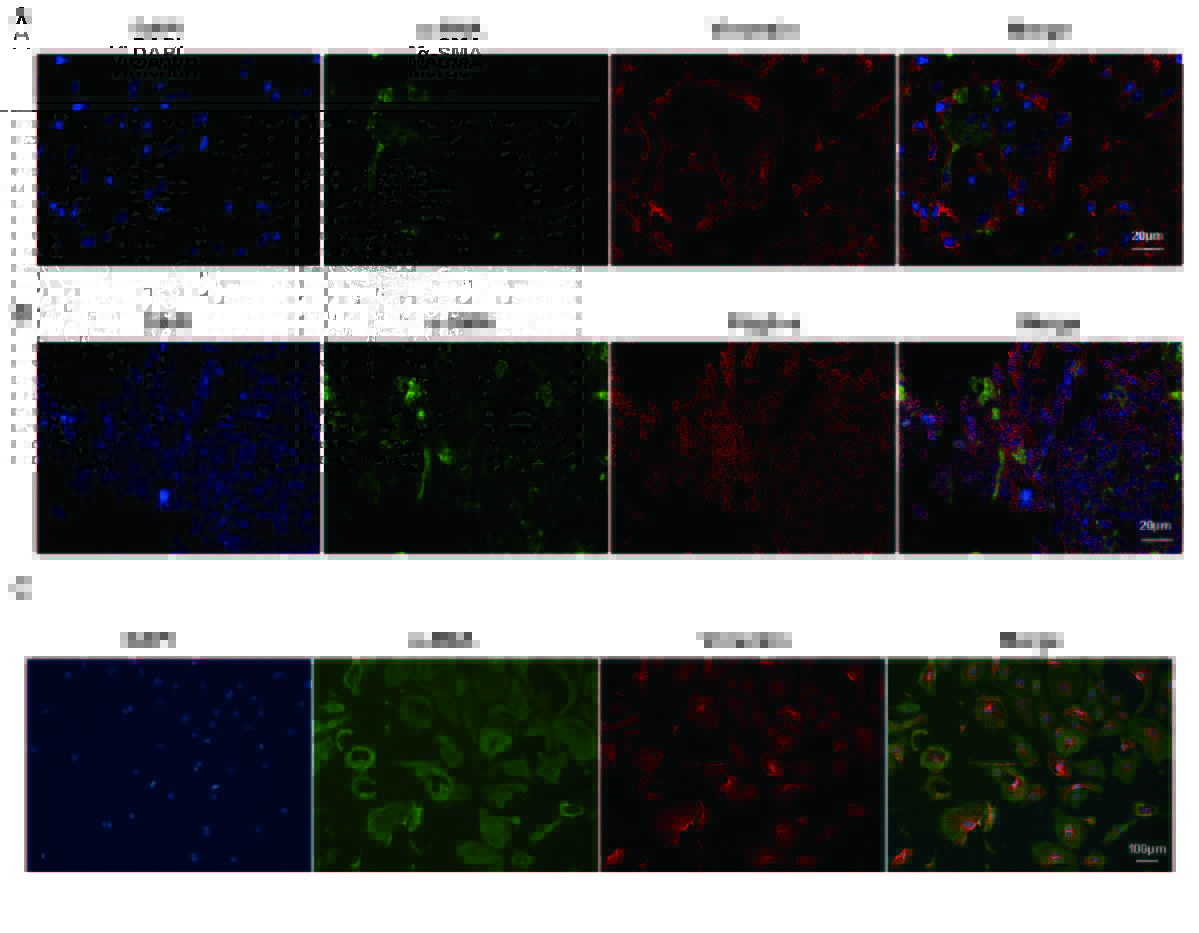

**Supplementary figure 5**. **Separate channels of immunochemistry.** (A) Separate channels of α-SMA and vimentin positive fibroblasts in explant. (B) Separate channels of α-SMA and Pdgfr-α positive fibroblasts in explant. (C) Separate channels of α-SMA positive primary fibroblasts on polyacrylamide substrate.
